# A unifying model for the propagation of prion proteins in yeast brings insight into the [*PSI*^+^] prion

**DOI:** 10.1101/2020.01.13.904060

**Authors:** Lemarre Paul, Sindi S. Suzanne, Pujo-Menjouet Laurent

**Affiliations:** Institut Camille Jordan, CNRS UMR 5208, University of Lyon, Villeurbanne, France; School of Natural Sciences, University of California Merced, Merced, CA, USA; INRIA Rhônes-Alpes, INRIA, Villeurbanne, France

## Abstract

The use of yeast systems to study the propagation of prions and amyloids has emerged as a crucial aspect of the global endeavor to understand those mechanisms. Yeast prion systems are intrinsically multi-scale: the molecular chemical processes are indeed coupled to the cellular processes of cell growth and division to influence phenotypical traits, observable at the scale of colonies. We introduce a novel modeling framework to tackle this difficulty using impulsive differential equations. We apply this approach to the [*PSI*^+^] yeast prion, which associated with the misconformation and aggregation of Sup35. We build a model that reproduces and unifies previously conflicting experimental observations on [*PSI*^+^] and thus sheds light onto characteristics of the intracellular molecular processes driving aggregate replication. In particular our model uncovers a kinetic barrier for aggregate replication at low densities, meaning the change between prion or prion-free phenotype is a bi-stable transition. This result is based on the study of prion curing experiments, as well as the phenomenon of colony sectoring, a phenotype which is often ignored in experimental assays and has never been modeled. Furthermore, our results provide further insight into the effect of guanidine hydrochloride (GdnHCl) on Sup35 aggregates. To qualitatively reproduce the GdnHCl curing experiment, aggregate replication must not be completely inhibited, which suggests the existence of a mechanism different than Hsp104-mediated fragmentation. Those results are promising for further development of the [*PSI*^+^] model, but also for extending the use of this novel framework to other yeast prion or amyloid systems.

**Author summary:** In the study of yeast prions, mathematical modeling is a powerful tool, in particular when it comes to facing the difficulties of multi-scale systems. In this study, we introduce a mathematical framework for investigating this problem in a unifying way. We focus on the yeast prion [*PSI*^+^] and present a simple molecular scheme for prion replication and a model of yeast budding. In order to qualitatively reproduce experiments, we need to introduce a non-linear mechanism in the molecular rates. This transforms the intracellular system into a bi-stable switch and allows for curing to occur, which is a crucial phenomenon for the study of yeast prions. To the best of our knowledge, no model in the literature includes such a mechanism, at least not explicitly. We also describe the GdnHCl curing experiment, and the propagon counting procedure. Reproducing this result requires challenging hypotheses that are commonly accepted, and our interpretation gives a new perspective on the concept of propagon. This study may be considered as a good example of how mathematical modeling can bring valuable insight into biological concepts and observations.

## Introduction

Amyloids are self-perpetuating protein aggregates, involved in many neurodegenerative diseases in mammals such as Alzheimer’s Disease, Parkinsons’s Disease, Creutzfeldt-Jakob Disease, Huntington’s Disease [1]. Prions are a particular case of amyloids, where a conformational change is infectious and transmissible through self-catalyzed aggregation [2]. Understanding the molecular mechanisms associated with amyloid growth and prion replication is crucial in the effort against neurodegenerative diseases. The yeast *Saccharomyces cerevisiae* has emerged as a powerful model system in the study of these processes [3]. Native yeast prions share many specificities with mammalian prions [4], and the protein quality control machinery involved in their propagation is highly conserved between mammals and yeast. Furthermore, yeast models are used to screen for anti-amyloid drugs through the artificial expression of mammalian proteins [5]. Many kinetic models of aggregate growth and nucleation have been proposed [6], but their validation using data from yeast colonies is challenging. Indeed, the propagation of yeast prions is a multi-scale system where molecular processes are coupled to cellular processes such as cell growth and division to produce effects that are observable at the phenotypical scale, *i.e.* at the scale of colonies. An illustration is proposed in Fig 1. Taking into account and relating those scales calls for specific modeling approaches, but to our knowledge no mathematical modeling work describes the whole system rigorously and in a controlled way. We propose a novel approach based on the use of impulsive differential equations. It is important to emphasize that this framework is versatile and can be used to study the evolution of any chemically active intracellular components through mass-action kinetics inside growing yeast colonies.

**Fig 1.**
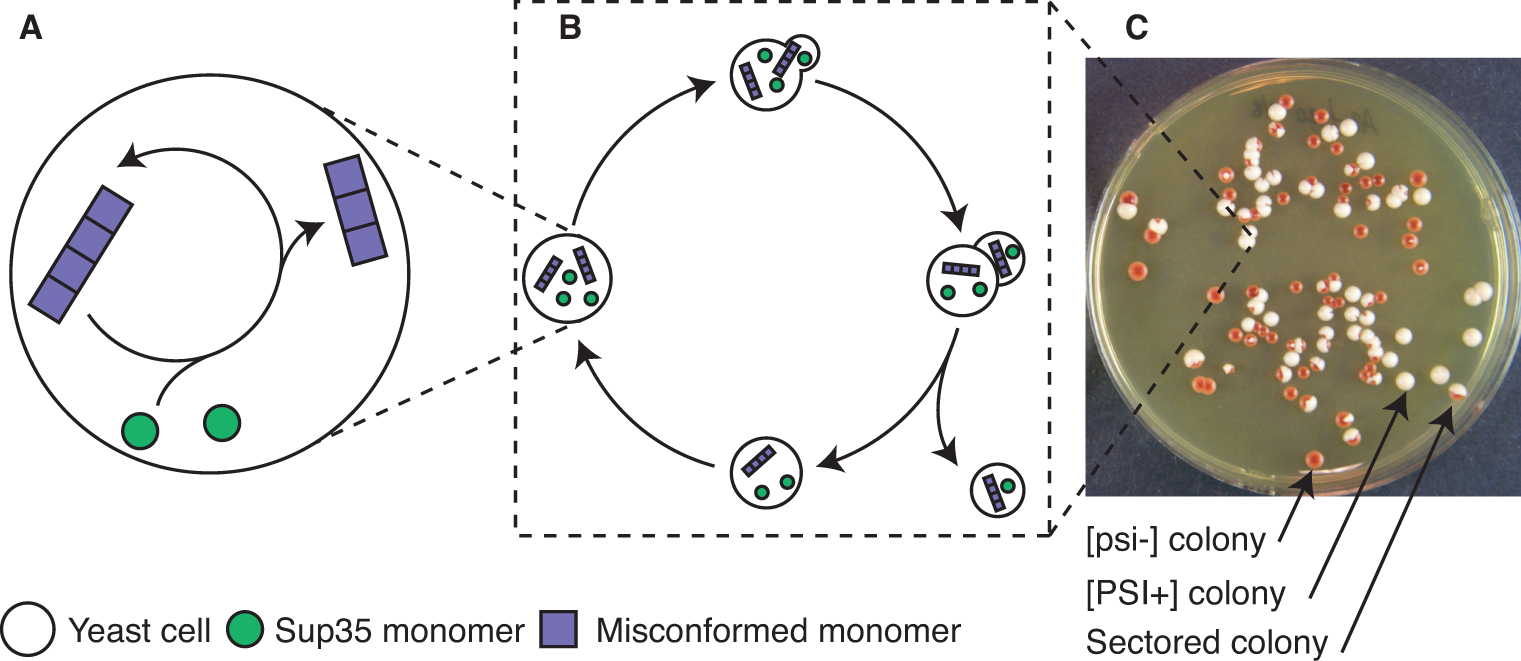
Different scales of yeast prion propagation. (A) Molecular processes of protein conversion and aggregation. (B) Growth and budding, along with aggregate transmission from mother to daughter. (C) Typical experimental observation of yeast colonies (circles) in a culture plate (courtesy of T. Serio). Each colony was founded by an isolated single cell.

As a first application of this approach, we focus here on the yeast prion [*PSI*^+^]. It has been studied in detail [7], and experimental results on [*PSI*^+^] have become essential groundwork for the development of prion biology [8, 9]. The [*PSI*^+^] phenotype is associated with the oligomerization and loss-of-function of the Sup35 protein, an enzyme release factor responsible that promotes stop-codon recognition. When aggregates are formed, a change of color of the yeast colonies is observed (from red to white) if they encode a stop-codon in either the ADE1 or ADE2 genes [10]. See Fig 1C for an illustration of the change in phenotype. As is the case with most native yeast prions, the [*PSI*^+^] phenotype is reversible. It can be cured using different physico-chemical treatments [11, 12] but also by introducing point mutations in Sup35 or its co-chaperones [13, 14]. Curing experiments are widely used to derive information on the molecular properties of Sup35 aggregates, depending on the curing treatment applied or the strain studied. The most exploited one is Guanidine Hydrochloride (GdnHCl) treatment, because it allows to infer prion seed numbers in [*PSI*^+^] cells [15, 16]. We reproduce some of those curing experiments using a model for aggregate replication at the molecular scale combined with a model for cell growth and division, as described in Figure 2. By coupling those aspects, we build a full model of prion propagation inside yeast colonies, which is detailed in Methods. Even by keeping things as simple as possible, our model sheds light onto two specific characteristics of the [*PSI*^+^] prion that were not uncovered in the past, as we detail in Results, the replication of aggregates at low densities follows non-linear effects, with a kinetic barrier preventing the expansion of the population of aggregates below a certain threshold. A second implication of our modeling work is that the effect of GdnHCl is not a complete block of aggregate fragmentation, as was suggested before [11], rather a significant reduction of the replication efficiency.

**Fig 2.**
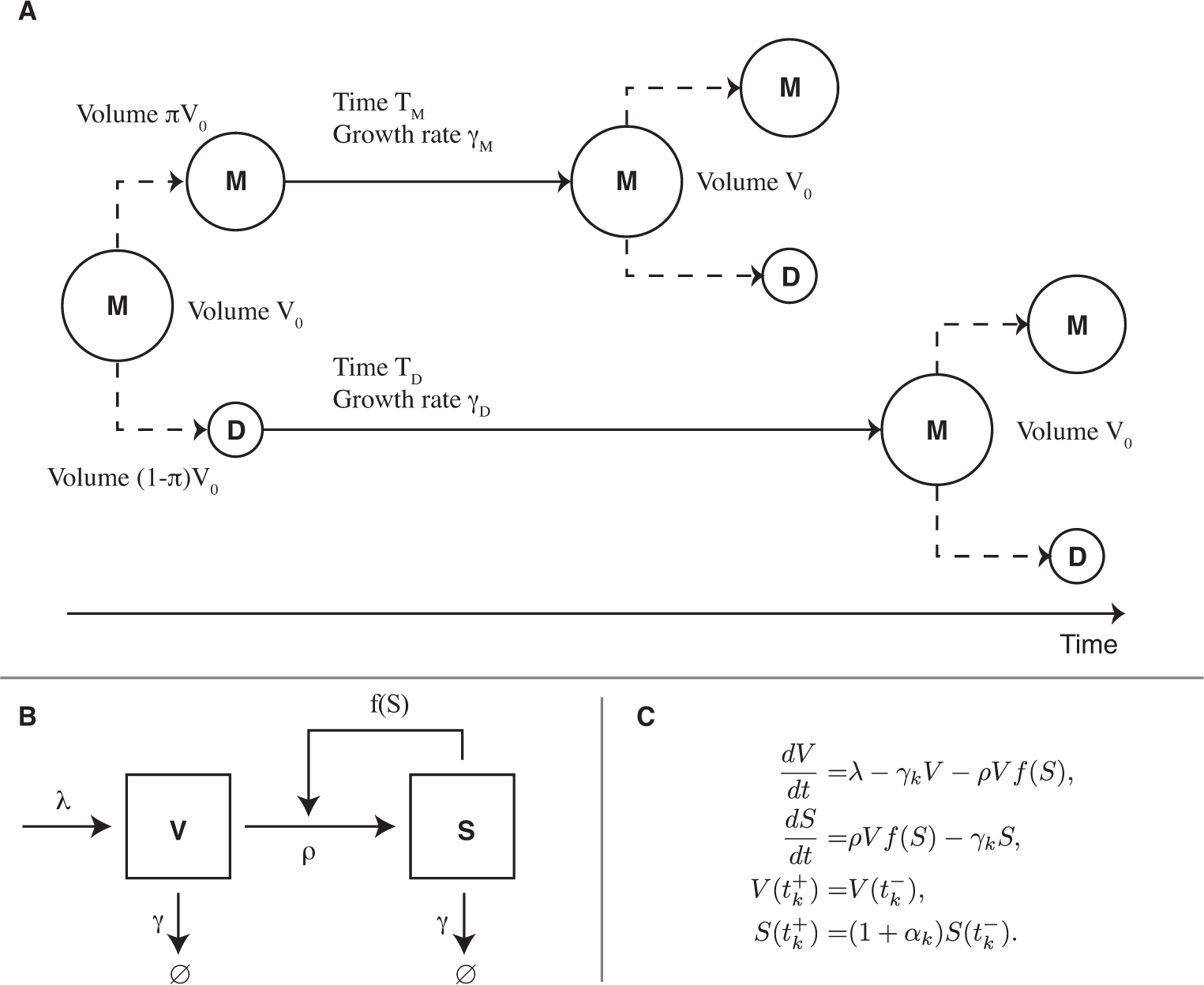
Multi-scale model of yeast prion propagation. (A) Cell growth and division model used for yeast budding. Cells are growing exponentially, and divide when they reach the volume *V*_0_. Division is asymmetrical and daughter cells have a smaller volume than mother cells. (B) Chemical model of aggregate replication, where *V* is the concentration of normal soluble Sup35 and *S* is the concentration of Sup35 in the prion conformation. (C) Corresponding equation system, the subscript *k* denotes the impulsion number. The function *f* is a sigmoid that accounts for non-linear cooperative effects of aggregate replication. See Methods for a description of each term.

## Results

We develop a multi-scale model of prion replication and propagation in yeast to investigate the relationship between molecular aggregation mechanisms and colony level phenotypical properties. The model is presented in Fig 2 and Table 1. A detailed mathematical introduction is led in Methods and an analytical study in S1 Appendix. In order to understand the results one should keep in mind that, in our model, newly born (daughter) yeast cells have a longer cell-cycle than their mothers. Furthermore, aggregate transmission at the moment of cell division is biased towards retention in the mother. Our goal was to develop the minimal molecular model required to reproduce experimental observations. In this work, we compare to observations for the [*PSI*^+^] prion, but it must be stressed that it could be adapted to any amyloid or prion propagating in growing yeast colonies. Using experimental results from the literature, we build a simple yet satisfying model and derive conclusions regarding the molecular mechanism of aggregate replication.

**Table 1.** Parameter values used by default in the simulations

| Parameter | Value | Unit | Description |
| --- | --- | --- | --- |
| $T_M$ | $2^*$ | hr | Mother doubling time |
| $T_D$ | $3^*$ | hr | Daughter doubling time |
| $\pi$ | $0.6^\dagger$ | - | Mother/daughter volume ratio |
| $\varepsilon$ | 0.1 | - | Mass transmission bias |
| $\alpha_M$ | $\frac{\varepsilon}{\pi}$ | - | Mother concentration bias |
| $\alpha_D$ | $-\frac{\varepsilon}{1-\pi}$ | - | Daughter concentration bias |
| $\gamma_M$ | $-\frac{1}{T_M} \ln(\pi) = 0.26$ | $\text{hr}^{-1}$ | Mother growth rate |
| $\gamma_D$ | $-\frac{1}{T_D} \ln(1 - \pi) = 0.31$ | $\text{hr}^{-1}$ | Daughter growth rate |
| $\lambda$ | $0.7^{\ddagger}$ | $\mu M \cdot \text{hr}^{-1}$ | Sup35 monomer basal production rate |
| $\rho$ | 10 | $\text{hr}^{-1}$ | Maximal aggregate replication rate |
| $K$ | 1 | $\mu M$ | Replication efficiency threshold |
| $n$ | 5 | - | Replication efficiency order |

### Aggregate replication is limited by a kinetic barrier

#### The [*PSI*^+^] phenotype is reversible

We investigate the general concept of [*PSI*^+^] curing. A cell is considered free of [*PSI*^+^] if, when plated onto regular medium (free of any treatment) it grows into a [*psi*^*−*^] colony, as determined by a color assay (*i.e.* it grows into a dark red colony). In the same exact environmental and genetic conditions cells can grow either into [*PSI*^+^] or [*psi*^*−*^] colonies, depending on the history of treatments they have sustained, as is illustrated in Fig 1C. This property is exploited to track the dynamics of curing during continuous treatment with a de-stabilizing agent [11, 12, 15]. In order to reproduce experimental results involving [*PSI*^+^] curing, this is an essential aspect that must be captured by the model.

#### A multi-stable molecular model

In terms of mathematical modeling, the fact that different outcomes are possible in the same conditions is interpreted as multi-stability. Different stable solutions are able to attract the trajectories of the system, and the outcome depends on the initial condition Our model introduces this property by including a non-linearity in the replication rate of aggregates, with a sigmoidal dependency on the concentration of aggregates portrayed by the function *f* in the model equations (see Fig 2). This shape of function is frequently used to model cooperative reactions [20], it appears naturally from ligand-receptor interaction equations. Without knowing the specific kinetic scheme, we empirically choose a Hill function. With this non-linearity, the colony grown from a cell will exhibit one of three possible behaviors depending only on its initial condition.

- A full [*PSI*^+^] (totally white) colony, when all cells have a positive concentration of aggregates.
- A full [*psi*^*−*^] (dark red) colony, when all cells have a zero concentration of aggregates.
- A sectored colony (part white, part red), when parts of the colony are [*PSI*^+^] and others are [*psi*^*−*^].

Numerically, the aggregate concentration never reaches exactly zero even when the solution is converging towards the prion-free solution. In practice, when needed, we differentiate between a [*PSI*^+^] cell and a [*psi*^*−*^] cell by comparing the aggregate concentration to a critical value, that was set to 0.5*µM*. Fig 3 shows an example of simulation output for each of those cases, and Fig 2C shows a typical experiment exhibiting all three possible cases of phenotypes. It must be emphasized that without the bi-stability introduced in the molecular kinetic scheme, those three outcomes would not be possible simultaneously (*i.e.* without changing the model parameters). This result is justified mathematically in Methods. These only represent the fate of three different cells growing in the same exact conditions, but starting with a different aggregated state of Sup35. Those three types of behavior are observed in every curing experiment monitored by color assays [7]. By numerically investigating the model, we derive a map linking the aggregated state of Sup35 in the founder cell to the phenotype of the colony grown from that cell, as shown in Fig 4. This map is numerically derived from the study of the two extreme lineages, mother-only and daughter-only. Those lineages correspond to periodic impulsive systems, and the corresponding solutions are attracted to periodic solutions. In each case (mother system or daughter system), two different periodic solutions are locally stable, a prion-free periodic solution and a [*PSI*^+^] periodic solution. A similar behavior was proven with a one-dimensional bi-stable equation in [21], but our results are only numerical so far. Those solutions are denoted on the phenotype map in Fig 4, corresponding to the first point in each periodic impulsion. They provide information regarding the whole colony, because they represent the most extreme behavior imposed by the transmission bias. After a sufficient amount of time, any lineage in the colony is predicted to be in-between those two lineages.

**Fig 3.**
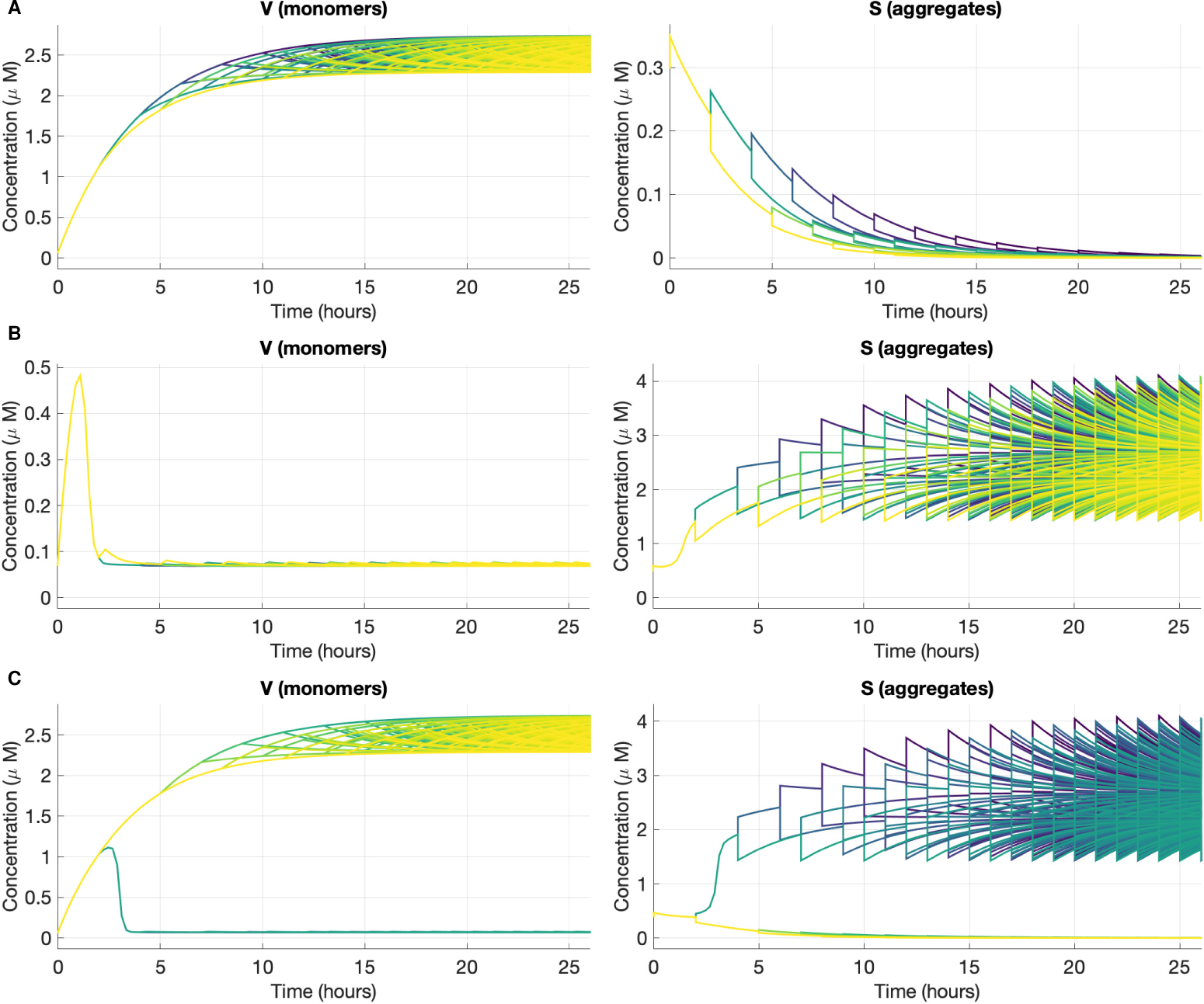
Representative simulation outputs of the bi-stable impulsive model. Evolution of the aggregate *S* and monomer *V* concentration for three different initial conditions. Each trajectory shown corresponds to a different lineage. Because cells are dividing the number of trajectories plotted increases in time, as does the number of cells. The trajectories for *S* and *V* are shown in the same color for the same cell. The parameters used are described in Table 1. The initial conditions used are to be related with the attraction basin showed in Figure 4. (A) Completely cured ([*psi*^*−*^]) colony (*V* (0) = 0.07*µM, S*(0) = 0.3*µM*) (B) Full [*PSI*^+^] colony (*V* (0) = 0.07*µM, S*(0) = 0.5*µM*) (C) Sectored colony (*V* (0) = 0.07*µM, S*(0) = 0.4*µM*).

**Fig 4.**
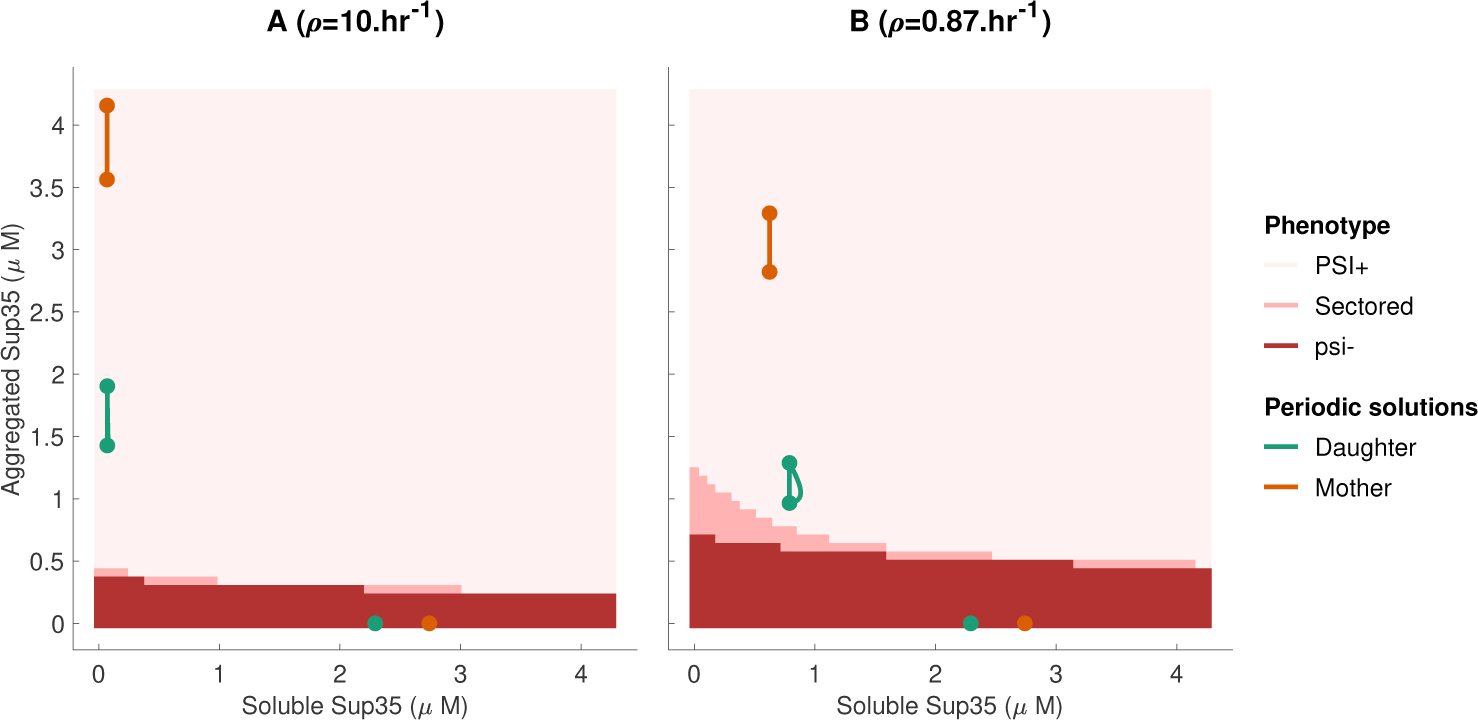
Predicting colony color phenotype. Numerical prediction of colony phenotype based on the state of the founder cell, for two different strains. The periodic solutions for the mother-only and daughter-only solutions are also shown. The parameters used for this diagram are described in Table 1. Panel (A) shows a strong strain with a maximal replication rate *ρ* = 10hr^*−*1^ and panel (B) is a weak strain with *ρ* = 0.87hr^*−*1^

#### The particular case of colony sectoring

The phenomenon of colony sectoring is the consequence of both the kinetic barrier and asymmetrical division. Indeed, our model includes a transmission bias that favors retention of aggregates by mother cells. (This bias is supported by numerous studies [7, 9, 22].) This means that under conditions of [*PSI*^+^] de-stabilization, daughter cells are more likely to fall under the threshold of efficient aggregate sustaining, and become [*psi*^*−*^]. This provides an explanation for sectoring that is relevant with many aspects of the biology. First, sectoring occurs more often when cells are treated with de-stabilizing agents. In Fig 4, we see how reducing the proportion of aggregated Sup35 will bring cells into the region corresponding to a sectored phenotype. Furthermore, we also know that “weak” strains are more likely to exhibit sectors [9].Fig 4 (B) shows the phenotype map for a strain that has a slightly reduced aggregate replication rate. We see immediately that the region leading to sectored phenotypes is enlarged, and also closer to the periodic solutions. This means that such a strain is more likely to be de-stabilized by random perturbations or by chemical treatment. In the literature, sectoring is often dismissed by counting mosaic colonies as either full [*PSI*^+^] or half and half [3, 11]. Perhaps some valuable information could be obtained by studying the phenomenon in more detail.

### GdnHCl does not completely block aggregate replication

#### The propagon count experiment

Among the various agents that de-stabilize [*PSI*^+^], GdnHCl is one of the most studied. It acts by impairing the chaperone Hsp104, which fragments Sup35 aggregates, thus preventing the creation of new aggregates [11]. [*PSI*^+^] colonies growing under GdnHCl treatment progressively lose the prion phenotype, by dilution of the aggregates between dividing cells. This property was used to infer the number of prion “seeds” in individual cells [15] by fitting an exponential model to the curing curve. This approach was supported by [3], with the discovery that during GdnHCl treatment, the number of [*PSI*^+^] revertants (cells that will grow into a [*PSI*^+^] colony when transferred back onto regular medium) reaches a plateau. This number was interpreted as the number of propagons in the initial cell, where propagon refers to the hypothetical prion elementary seed. The interpretation of the GdnHCl curing experiment is illustrated by Fig 5. Propagon counts and statistics on those counts are often used to characterize strains [23], to quantify the bias of transmission between mother and daughter cells [17], to study the effect of different point mutations [13]. It needs to be emphasized that this interpretation is based on two hypotheses. The first hypothesis is that the threshold between a [*PSI*^+^] cell and a [*psi*^*−*^] cell is the presence of aggregates or not. In our modeling framework, this hypothesis translates to the presence of a concentration threshold under which aggregate replication is inefficient. The second hypothesis is that GdnHCl instantaneously and totally blocks the creation of new aggregates. Our results contradict this second hypothesis as explained below.

**Fig 5.**
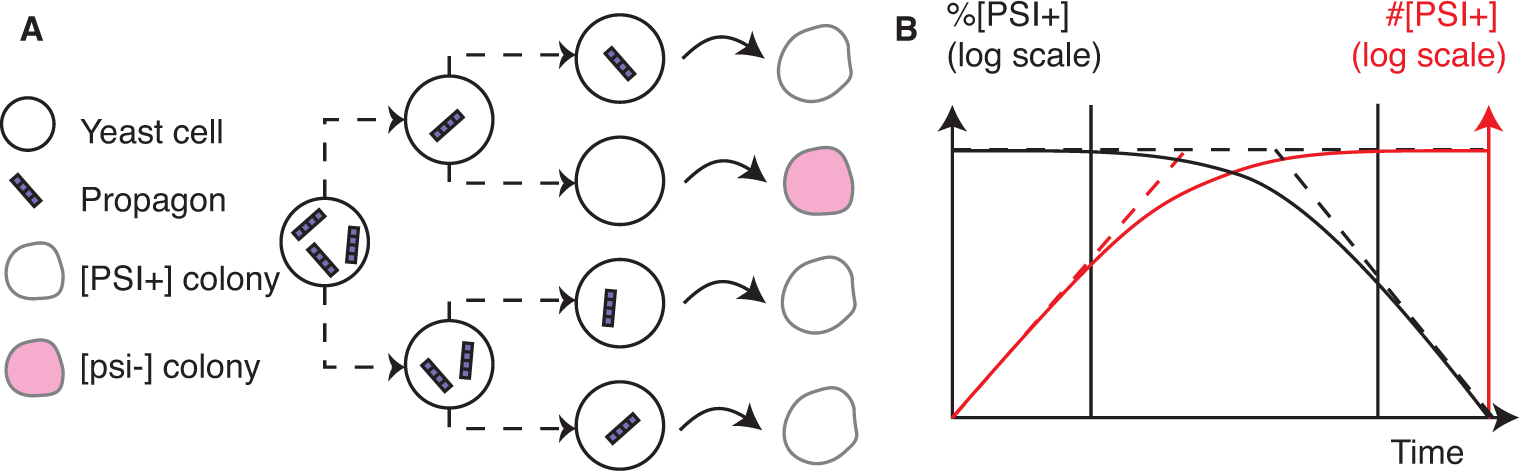
The GdnHCl curing experiment and propagon counting. (A) Two phase propagon counting experiment. Propagons are distributed in dividing cells under GdnHCl treatment. Cells are then plated onto regular medium and the color phenotype ([*PSI*^+^] or [*psi*^*−*^]) of each subsequent colony is determined by color assay. The number of [*PSI*^+^] colonies is counted and gives the propagon count. (B) Curves obtained during the GdnHCl curing experiment. In black is the percentage of [*PSI*^+^] cells in the colony (logarithmic scale), in red is the number of [*PSI*^+^] cells in the colony (logarithmic scale). The plateau in the red curve corresponds to the propagon count. See [9] for an experimental example of such curves.

#### The propagon plateau

During GdnHCl treatment, the number of [*PSI*^+^] cells in a colony reaches a plateau [9]. Our model reproduces this property under specific parameter choices, as shown in Fig 6. In those simulations, cells are scored as [*PSI*^+^] as soon as the concentration of aggregated Sup35 is above a given threshold (we used 0.5*µM*). However the qualitative result does not depend on the scoring method, because we know that a finite number of lineages in the colony are attracted by a solution with a positive concentration of aggregates. All the other lineages are attracted by the prion-free solution. This behavior is possible with our model but only if aggregates continue to replicate in GdnHCl conditions, which strongly contradicts former experimental studies [11, 15]. The hypothesis that GdnHCl interrupts all chemical activity of aggregates was nuanced in previous experimental work. Indeed it was shown GdnHCl does not stop aggregate growth [11, 17, 24]. Furthermore, [25] used fluorescent tagging to track aggregates during GdnHCl treatment, and report a decrease in cells with foci slower than the halving predicted by the segregation hypothesis.

**Fig 6.**
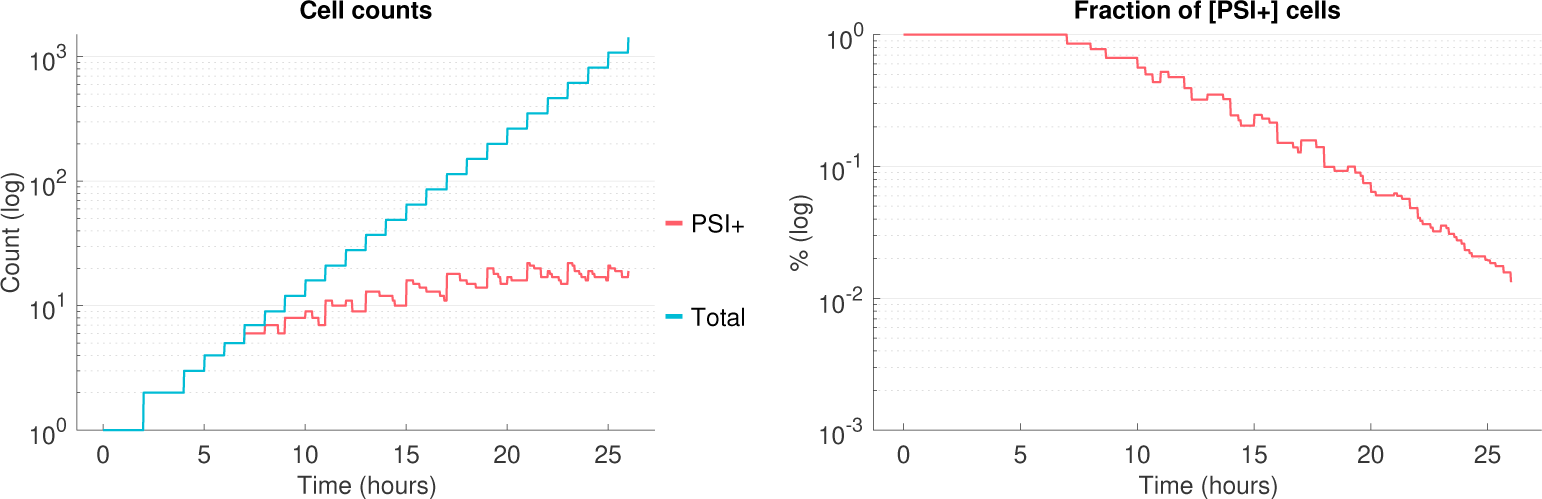
Reproducing a propagon count experiment. Evolution of the [*PSI*^+^] cells count and [*PSI*^+^] proportion simulated by our model, in the case of *ρ* = 0.21hr^*−*1^ (all other parameters as in Table 1). Cells are scored as [*PSI*^+^] as soon as they contain a concentration of aggregated Sup35 higher than 0.5*µM*.

Reproducing the propagon count experiment with our model requires very specific kinetic parameters for the replication reaction. Those conditions are found numerically by studying the mother-only lineage in the model. Indeed when this lineage is attracted to a [*PSI*^+^] solution but each of its daughters becomes [*psi*^*−*^], we are assured that the total number of [*PSI*^+^] cells in the colony reaches a plateau. Once again this is the consequence of bi-stability in the molecular model as well as the asymmetric division favoring aggregate retention by mother cells. The fact that those conditions correspond to a very narrow parameter range is worth emphasizing and discussing further.

## Discussion

### How to explain the concentration threshold for aggregate replication

In previous modeling work, one approach was to study aggregates as discrete entities [15, 26, 27]. When doing so, the kinetic barrier hypothesis is implicit and the threshold lies at the difference between 0 and 1 aggregate. However, it is not a proper modeling approach when it comes to studying chemical reactions, because mass-action kinetics cannot be applied to numbers in isolation but rather to densities. Our model implements chemical modeling inside a multi-scale phenomenon in a rigorous way and provides a powerful tool for generating insights into existing experimental outcomes: colony sectoring and propagon counting assays.

Our work suggests that the multi-stability in the intracellular scale is essential to reproduce the experimental results for yeast prions. That is, the prion-free state and the prion state both need to be locally stable. But what is the mechanistic origin of this multi-stability? The non-linearity we introduced could be interpreted by a cooperativity phenomenon, where the replication of aggregates (for instance fragmentation) is chemical reaction that is catalyzed by the presence of other aggregates. Due to the lack of clear biological evidence, there is so far no real hypothesis to explain such a mechanism. One idea is to take into account the action of chaperones, in particular Hsp104. However, even if fragmentation is limited by the presence of Hsp104 the model does not exhibit bi-stability [28]. We would like to emphasize that this type of modeling (global models) is universally used in the prion biology, in particular with the widespread use of the Nucleated Polymerization Model (NPM) [29, 30]. The introduction of bi-stable systems was suggested in earlier work [31, 32], in the case of mammals and more recently to study interaction between prion strains [33]. The benefit of the framework we suggest here is that it could allow to use data to validate a precise kinetic scheme of aggregate replication. In particular, the sectoring phenomenon should be studied with a new perspective, especially during curing experiments.

### What is the effect of GdnHCl?

Our results question all previous assumptions made about the effect of GdnHCl. With our model, the propagon experiment is reproduced only if the chemical replication rates are chosen very carefully. This is potentially explained by two reasons. The first is that our model might be too simple to capture the possibilities of the full biological system. This would also explain why sectoring is reduced to such a narrow region in the phenotype map Fig 4.

A second explanation is that our model may be missing an essential chemical process, which remains unaffected when GdnHCl is present. There is precedent for this assumption, because there is evidence that aggregates are still chemically active under GdnHCl treatment. First GdnHCl does not prevent aggregates from growing by polymerizing newly synthesized Sup35 [17, 34]. Furthermore, there is evidence for an action of Hsp104 that is not affected by GdnHCl [35, 36]. This brings up the controversial question of the roles of Hsp104 in the propagation of [*PSI*^+^] and other yeast prions. The only conclusion our results bring is that GdnHCl curing is not explained by an exponential dilution model as suggested previously [15, 18], because of the plateau of [*PSI*^+^] cells. Our study is a first step in the design of a more elaborate kinetic model that includes the effect of Hsp104.

## Conclusion and Perspectives

We introduced a novel modeling tool to the field of yeast prions, with the major benefit of relating different scales in a controlled and rigorous way. From the molecular mechanisms, with a kinetic scheme built using mass-action kinetics, to the phenotypical traits at the colony level, this framework has the potential of taking into account every aspect of the system. In the case of the [*PSI*^+^] prion, we build a simple model with the primary goal to qualitatively reproduce experimental observations from the literature. We focus on curing experiments, and reproducing any curing experiment with our framework requires introducing a very particular characteristic into the molecular model. The kinetic scheme needs to be bi-stable, where the prion-free state and the prion state are both simultaneously stable, and the transition between the two of them is a bi-stable switch. This allows the possibility of curing, and it concomitantly explains the phenomenon of sectoring in a deterministic way. This phenomenon is often dismissed but is in fact instructive with regards to the molecular processes. By investigating in more detail the case of GdnHCl curing, we have reason to question the suggested effect of this agent. Indeed this experiment is reproduced by our model, but the fragmentation of aggregates must not be completely inhibited contrary to the commonly accepted effect of GdnHCl.

Overall, the framework of impulsive differential equations is versatile and could be adapted to many different cases. Studying the [*PSI*^+^] prion already revealed instructive, even though work is still in progress. In the future, we aim to use this framework to build and validate a complete model of aggregate replication including the role of Hsp104 and possibly its co-chaperones, a size-distribution of aggregates, stochasticity in the cell division events. This would be done through hypothesis testing and close collaboration with biologists. Inferring parameters is a long-term goal, that first requires understanding the very structure of the molecular processes. In particular, it needs to be clear what is the mechanistic origin of the cooperativity in the replication of aggregates. Another use of the model is to extend it to other yeast prions and amyloid models, as yeast models are used to screen for anti-amyloid drugs, and a specific modeling framework would help interpreting those experiments.

## Methods

### Using impulsive differential equations to model cell division

We first introduce impulsive differential equations (IDEs) and use them to model a population of dividing yeast cells. Given a system of ordinary differential equations *f* (*Z, t*), where *Z* ∈ ℝ^*n*^ represents the concentrations of the biochemical species we are tracking, and given *Z*_0_ ∈ ℝ^*n*^, *Z*_0_ ≥ 0 an initial condition, we define the impulsive system

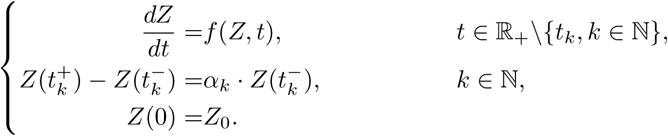

The sequence 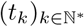 represents the impulsion times, which we choose to be fixed. We require 0 ≤ *t*_0_ < … < *t*_*k*_ … and lim_*k*→∞_ *t*_*k*_ = *∞*. Each impulsion corresponds to a cell division, and the vector *α*_*k*_ ∈ℝ^*n*^ represents the partition rate at the moment of division *k*. A solution to this system is a piece-wise continuous function *Z* on (*t*_*k*_, *t*_*k*+1_], for 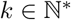. The existence and the properties of such solutions are studied theoretically [37, 38]. IDEs are used in various modeling fields, including epidemiology [39], population dynamics [40] and more recently T-cell differentiation [21].

In order to model cell division with IDEs, we need to clarify how species are distributed between mother and daughter cells. Consider a yeast cell about to bud and produce a daughter. The volume of the mother cell is *V*_0_, which will split into two parts right after division. The mother’s volume becomes *V*_*M*_ = *πV*_0_ and the daughter gets *V*_*D*_ = (1 *− π*)*V*_0_, where *π* is the volume asymmetry. For yeast cells it has been observed that *π* = 0.6 [18]. Next consider a chemical component in the mother of mass *M*_0_ and concentration 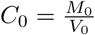. During division this mass is distributed between the mother and the daughter cells. With no bias (fast diffusion) it should distribute with the same ratio as the volume. If we introduce a bias, the mother retains the mass *M*_*M*_ = (*π* + *ε*)*M*_0_ and the daughter receives *M*_*D*_ = (1 − *π* − *ε*)*M*_0_. The parameter *ε* represents the bias, with a positive value corresponding to a retention of the mass by the mother. In order to ensure positive mass we require

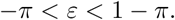

After division, the mother concentration (*C*_*M*_) and the daughter concentration (*C*_*D*_) are given by

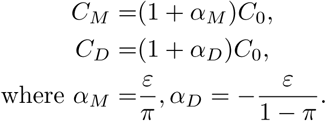

We dot not consider degenerate cases where *ε* = −*π* or *ε* = 1 − *π*, corresponding to the case where the transmission from mother to daughter is (respectively) full or null. This hypothesis is biologically relevant in the sense that it seems highly unlikely that exactly no misconformed protein be transmitted at the moment of division.

We assume that cells are exponentially growing, and that they divide when they reach the volume *V*_0_. In other words, the intracellular dynamics of prion aggregation have no impact on the division cycle. This is consistent with [17]. This assumption imposes a relation on the growth rate *γ* and the division time *T*. As before, if we denote the mother parameters with a subscript *M* and the daughter parameters with a subscript *D*, we have the following two relations

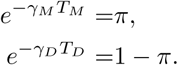

As a consequence when *π*, *T*_*M*_ and *T*_*D*_ are fixed (as measured experimentally [9, 17]), *γ*_*M*_ and *γ*_*D*_ are determined by these relations. Figure 2A shows a visual representation of the cell division model we are using. (The parameters we used in results are given in Table 1.)

An impulsive differential equation model allows one to follow a cell lineage and its intracellular components in time. The cell history is described by the sequence of impulsions it undergoes, either mother or daughter impulsions. Analytically, we study in detail the two extreme lineages, mother-only and daughter-only. Indeed those lineages are subject to periodic and identical impulsions, which facilitates the analysis [37]. If we use the IDEs to track all lineages starting from a single cell, we can study the complete yeast colony. Having formalized the cell division dynamics model, we next introduce the model for intracellular dynamics.

### A bi-stable prion replication process

In this section, we introduce a two-dimensional bi-stable model of prion dynamics, at first without considering the effect of impulsions. In each yeast cell, we track the concentration of soluble Sup35 (*V*), and of Sup35 in the prion conformation (*S*). The model is illustrated in Figure 2B, and described by the following system of ordinary differential equations

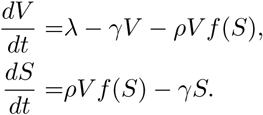

The parameters *λ* and *γ* are respectively the Sup35 monomer production rate and the cell growth rate. For simplicity we do not include the aggregate size dynamics but model the recruitment of monomers into prion aggregates. It occurs with a cooperative reaction, at maximal rate *ρ* and non-linear efficiency *f* (*S*). The function *f* is chosen to be a Hill function of order *n*(*>* 1) and threshold *K*

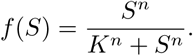

This function is chosen for one distinct feature: it makes the system multi-stable as soon as *n >* 1. The prion-free equilibrium (with *S* = 0) exists under any choice of parameters and remains locally stable. Two other equilibria appear from a saddle-node bifurcation, one of them is unstable and the other is locally stable. In conditions when the three equilibria exist (one prion-free and two prion equilibria), the asymptotic outcome of the model depends on the initial condition. Starting with a concentration of aggregates too low will cause the solution to converge to the prion-free equilibrium, whereas if the initial concentration of aggregates is sufficient their population will be stably maintained. See S1 Appendix for the mathematical description and proof of this property, as well as some numerical illustration of the bi-stability. As detailed in Discussion, it is essential that we use a model with multi-stability because we are investigating a curing experiment, where cells can lose the prion phenotype.

### Complete bi-stable model with impulsions

When combining the bi-stable model and the impulsive differential equation framework, we obtain a complete model of aggregate replication and transmission in growing yeast colonies. The choice of parameters used by default is detailed in Table 1. The cell division parameters (doubling time and mother-daughter volume ratio) are measured experimentally and we choose values within the typical observed range [16, 17]. The mass transmission bias is a parameter that will require a thorough investigation in further work but we choose a reasonable value of *ε* = 0.1 for the time being. The Sup35 monomer production rate is such that the prion-free steady-state concentration of Sup35 in cells is 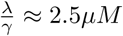[19]. The prion replication parameters *ρ*, *K* and *n* are used as adjustment parameters to investigate the behavior of the model.

## Supporting information

**S1 Appendix. Analytical study of the bi-stable impulsive system for yeast prion propagation.**

## Acknowledgments

The authors would like to thank Dr. Tricia R. Serio for a contributing a figure of yeast colonies and offering valuable comments.

